# Transcriptomics provides a robust framework for the relationships of the major clades of cladobranch sea slugs (Mollusca, Gastropoda, Heterobranchia), but fails to resolve the position of the enigmatic genus *Embletonia*

**DOI:** 10.1101/2020.09.22.307728

**Authors:** Dario Karmeinski, Karen Meusemann, Jessica A. Goodheart, Michael Schroedl, Alexander Martynov, Tatiana Korshunova, Heike Wägele, Alexander Donath

## Abstract

**Background:** Cladobranch sea slugs represent roughly half of the biodiversity of soft-bodied, marine gastropod molluscs (Nudibranchia) on the planet. Despite their global distribution from shallow waters to the deep sea, from tropical into polar seas, and their important role in marine ecosystems and for humans (as bioindicators and providers of medical drug leads), the evolutionary history of cladobranch sea slugs is not yet fully understood. Here, we amplify the current knowledge on the phylogenetic relationships by extending the cladobranch and outgroup taxon sampling using transcriptome data.

**Results:** We generated new transcriptome data for 19 species of cladobranch sea slugs and two additional outgroup taxa. We complemented our taxon sampling with previously published transcriptome data, resulting in a final supermatrix covering 56 species from all but one accepted cladobranch superfamilies. Transcriptome assembly using six different assemblers, selection of those assemblies providing the largest amount of potentially phylogenetically informative sites, and quality-driven compilation of data sets resulted in three different supermatrices: one with a full coverage of genes per species (446 single-copy protein-coding genes) and two with a less stringent coverage (667 genes with 98.9% partition coverage and 1,767 genes with 86% partition coverage, respectively). We used these supermatrices to infer statistically robust maximum-likelihood trees. All analyses, irrespective of the data set, indicate maximum statistical support for all major splits and phylogenetic relationships on family level. The only discordance between the inferred trees is the position of *Embletonia pulchra*. Extensive testing using Four-cluster Likelihood Mapping, Approximately Unbiased tests, and Quartet Scores revealed that its position is not due to any informative phylogenetic signal, but caused by confounding signal.

**Conclusions:** Our data matrices and the inferred trees inferred can serve as a solid foundation for future work on the taxonomy and evolutionary history of Cladobranchia. The correct placement of *E. pulchra*, however, proves challenging, even with large data sets. Moreover, quartet mapping shows that confounding signal present in the data is sufficient to explain the inferred position of *E. pulchra*, again leaving its phylogenetic position as an enigma.

## Background

Marine Heterobranchia (Gastropoda) have become a major focus as bioindicators to monitor the health of coral reefs [1–7]. They mainly prey on a high variety of marine sessile organisms, from algae to sponges, cnidarians, bryozoans and tunicates, and very often take up the chemical compounds of the food for their own defence. These “stolen” compounds have become of high interest for pharmacists in finding new drug leads for medical applications [8–10]. However, they are also of high interest in understanding the evolution of photosymbiosis and the role of “stolen” chloroplasts or even whole algal cells incorporated in the slugs’ body, which help the slugs survive starving periods or otherwise increase fitness [11–14]. Within marine Heterobranchia, the shell-less Nudibranchia have developed a variety of biological strategies that make them unique within Metazoa. Of particular interest is the sequestration of cnidocysts from the cnidarian prey, storing them in special morphological structures (cnidosacs) in exposed body areas, and the ability to mature the stolen cnidocysts (cleptocnides) in the cnidosac [15–18]. This unique defence system seems to have evolved only in one of the major nudibranch clades, the Cladobranchia, within which there are likely two independent origins [18].

Nudibranchia, with the two clades Cladobranchia and Anthobranchia, form a monophyletic group that is well explained by morphological features [19]. Recently, the sister group relationship to Pleurobranchomorpha (Pleurobranchida) as well as monophyly of Nudibranchia was confirmed by transcriptomic data [20]. The monophyly of Nudibranchia has also been confirmed in various molecular analyses using larger taxon sets, albeit small gene sets (see review in [21]). However, few studies have used both morphological and molecular methods to obtain and explain phylogenetic relationships within Cladobranchia. A comprehensive study of Anthobranchia (Doridida) applying both molecular phylogenetic and ontogenetic data was published recently [22]. Similar studies are still lacking for Cladobranchia.

Pola and Gosliner [23] tried to resolve the phylogeny of Cladobranchia using one nuclear and two mitochondrial genes: the study resulted in a topology that primarily consisted of an unresolved comb. Bleidissel [24] analysed the Aeolidida within the Cladobranchia, based on three genes (18S, 16S, and CO1), in order to investigate the evolution of the incorporation of algae from the genus *Symbiodinium* in certain sea slugs. In this study, for the first time, the paraphyly of the aeolidid family Facelinidae was shown. Similar to morphological data, the success of retrieving more reliable relationships based on few molecular markers increases, when working on family level. Recently, by the inclusion of the type species of the genus *Facelina*, the “true” family Facelinidae was revealed and the name Myrrhinidae resurrected for the second “facelinid” clade [25]. Korshunova and colleagues [26] studied the relationships within the former Flabellinidae, including representatives of many Aeolidida. The authors provided much evidence for the paraphyly of the former Flabellinidae, which they then split into five different families.

Recent analyses, using a large transcriptomic data set, provided the first robust cladobranch tree that enabled the study of evolution of food preferences [27, 28]. In a subsequent study, a broader data set with nearly 90 taxa was used to examine the evolution of the cnidosac [18], which is the main defence system of Aeolidida [29]. Similar defence structures have evolved independently in *Hancockia* [15], a genus assigned to Dendronotida [18]. However, the authors based their interpretations on a phylogenetic tree with partly low statistical support. Moreover, a few taxa showed relatively long branches compared to other members of the family (*Cerberilla*) or even the same genus (*Janolus*). Therefore, bias due to possible long branch artefacts cannot be excluded. A reduced data set was used by Goodheart and Wägele [30] to study the taxonomic relationship of an enigmatic pelagic cladobranch, the genus *Phylliroe*, to analyse morphological traits enabling a shift from a benthic life style into a pelagic form. With this study presented here, using an extended data set including 40 publicly available transcriptomes and combining them with 21 newly sequenced transcriptomes, we provide robust support for yet unresolved relationships and reconsider the phylogenetic position of the genus *Embletonia*, which has been assigned to various groups in the past without any current consensus [16, 18, 24, 31, 32]. Robustly resolved and reliably inferred phylogenetic trees that are not affected by confounding signal, but driven by “true” phylogenetic signal, are one prerequisite for answering questions about the evolutionary history of taxa and biological phenomena, such as the aforementioned evolution of the cnidosac and photosymbiosis. Therefore, only trees that reflect most likely the “true” history of species allow the inference of biological traits to understand biodiversity and its origin. Inferred trees resulting from methodological or computational inadequacy can lead to erroneous hypotheses (see, e.g., [33]). Taxa that diversified quickly and/or underwent rapid radiation events within a short period of time are especially difficult to analyse (see, e.g., [34] and several examples in [35]). Rapid radiation might also be the reason why for some marine Heterobranchia it seems so difficult to reliably infer a species tree [21, 23, 36, 37].

In order to obtain a statistically highly supported tree and to check whether ambiguous splits in this tree might be based on confounding and thus erroneous signal, we performed a thorough study on 57 cladobranch and four outgroup transcriptomes. A comparison of the results of various *de novo* transcriptome assemblers allowed us to specifically select those assemblies that showed the highest sequence coverage with respect to a reference orthologue set. After accounting for possible influences on phylogenetic inference, e.g., among-lineage heterogeneity and rejecting stationary, homogenous and time-reversible conditions we compiled three final data sets including 56 out of originally 61 species (discarding three species with a low coverage of the orthologue set as well as two species due to model violation (see Methods): 1) the full data set allowing gene partitions to be missing for single species, 2) a smaller intermediate data set in terms of the number of genes, but with less missing data, and 3) a strict data set only including gene partitions for which all species were present. In addition to careful preparation and processing of the data throughout all steps of the analyses, i.e. evaluating the most appropriate assemblies, identifying single-copy protein-coding orthologs, a thorough check of multiple sequence alignments, and optimization and evaluation of the final data sets, we comprehensively examined the ambiguously inferred position of *Embletonia* for alternative topologies with approximately unbiased (AU) tests [38], Four-cluster Likelihood Mapping [39, 40], and quartet puzzling [41] approaches.

## Results and Discussion

### Data preparation prior to phylogenetic analyses

A list with details on the 21 species with newly sequenced transcriptome data is provided in Supplementary Table S1, Additional File 2. Accession numbers for all species are given in Supplementary Table S2, Additional File 2.

#### Transcriptome sequencing and data processing

Paired-end sequencing resulted in approximately 7.5 Gbases of raw data per sample. For the newly generated transcriptomes, the number of complete read pairs ranged from 20,266,817 in *Calmella cavolini* to 43,524,035 in *Facelina rubrovittata* with a median of 24,882,673 (*Hancockia* cf. *uncinata*). After trimming of possible adapter sequences and sequence regions of low quality, the average read length of complete read pairs ranged from 118.1 bp in *Hermissenda emurai* to 139.6 bp in *Doto* sp. with a median of 133.8 bp in *Polycera quadrilineata* (Supplementary Table S3, Additional File 2). Details on sequence processing is provided in the Supplementary Text, Additional File 1. Transcriptome assembly using six different *de novo* assemblers per data set resulted in a total number of 366 assemblies, i.e. six assemblies for each of the 61 transcriptomic data sets (see Supplementary Text, Additional File 1 and Supplementary Table S4, Additional File 2).

#### Evaluation of transcriptome assemblies, orthology prediction, and alignment procedures

Evaluation of assembled transcriptomes and subsequently applying BUSCO version 3.0.0 [42] with the Metazoa set including 978 orthologs revealed a median of 731 (75%) complete BUSCO genes per sample (maximum: 943 complete BUSCO genes, fragmented: 27, missing: 8 in *Caloria elegans* assembled with BinPacker; minimum: 158 complete BUSCO genes, fragmented: 123 missing: 697 in *Doris kerguelenensis* assembled with BinPacker). All quality assessment results of the transcriptomes using BUSCO are summarised in Supplementary Table S5, Additional File 2.

We additionally evaluated the quality of all transcriptomes separately for each assembly method based on the results of orthology prediction and identified single-copy protein-coding genes with our custom-made orthologue set comprising 1,992 orthologues (see Methods and Supplementary Text, Additional File 1). Results were ranked based on the cumulative length of transcripts that were successfully assigned to the reference genes used to identify single-copy orthologues (OGs) in the transcriptomes (see Supplementary Table S6, Additional File 2). The cumulative lengths ranged from 82,409 bp in *Pseudobornella orientalis* (the genus was recently resurrected by Korshunova and colleagues [43]) (IDBA-Tran, 458 genes successfully assigned) to 784,043 bp in *Caloria elegans* (Shannon, 1,904 genes successfully assigned). The median was 472,305 bp for the cumulative length and 1,577 for the number of successfully assigned genes. The best assembly (according to the largest cumulative length) out of the six available per sample was selected as the representative transcriptome for the respective species. This transcriptome was used for all further downstream analyses and submitted to NCBI (see Supplementary Tables S2 and S7, Additional File 2). In order to reduce the amount of missing data in subsequent analyses we excluded three samples for which less than 60% of OGs included in the search had been identified: *Pseudobornella orientalis, Dermatobranchus* sp., and *Tritoniopsis frydis*. Furthermore, we only kept OGs for which at least 50% of the investigated 58 species had a positive hit. This resulted in 1,767 OGs that we subsequently used to generate multiple sequence alignments (MSAs) on amino acid level. Checking the MSAs for outlier sequences (i.e. putatively misaligned or misassigned amino acid sequences), we identified 897 sequences in 112 MSAs that were subsequently removed. Outliers were found in sequences from all remaining 58 species with the highest number of 30 outlier sequences in *Limenandra confusa* and the lowest number of eight outlier sequences in *Doris kerguelenensis* (median: 15 outliers, all details are provided in the Supplementary Text, Additional File 1 and Supplementary Table S8, Additional File 2).

Alignment masking resulted in masking of alignment sites in 1,519 out of 1,767 genes (Supplementary Text, Additional File 1) leaving ~ 71% of aligned unmasked sites for subsequent analyses.

#### Compilation, evaluation and optimization of data sets

Analysing the concatenated supermatrix using MARE v. 1.2-rc [44], AliStat v. 1.6 [45] for information content and data coverage, and SymTest v. 2.0.47 [46] for putative violating of stationary, (time-)reversible and homogenous (SRH) model conditions [47, 48] using the implemented Bowker’s matched pairs test of symmetry [49] led to the results shown in Supplementary Figures S1 and S2, Additional File 3.

With respect to the amount and distribution of missing data we initially compiled two data sets as described in the methods section. The data set allowing for the highest amount of missing data, termed “original unreduced data set”, was not further reduced after concatenation and comprised 58 species, 771,739 aligned amino acid positions and 1,767 gene partitions. The second data set with a full gene coverage for all 58 species (termed “original reduced data set”) comprised 143,859 aligned amino acid positions and 364 gene partitions. Analysing both data sets for violation of SRH model conditions with SymTest revealed that two species strongly violated the SRH conditions: *Calmella cavolini* and *Doris kerguelenensis* (Supplementary Figure S2, Additional File 3). The latter transcriptome, which likely belongs to an *Architectonia* species (personal communication Vanessa Knutson), was probably mislabeled in the repository from which it was downloaded. Therefore, the sequences belonging to these two species were removed entirely from all MSAs from the original unreduced data set. This newly created data set (termed “unreduced data set”) spanned a superalignment length of 771,706 amino acid positions including 1,767 gene partitions.

To reduce the amount of missing data, we compiled an “intermediate” data set featuring only those gene partitions for which at least one representative of the selected taxa was present (see Materials and methods, Supplementary Text, Additional File 1, and Supplementary Table S9, Additional File 2). This data set (termed “intermediate data set”) spanned a superalignment length of 271,732 amino acid positions and included 667 gene partitions. The third and most strict data set with full gene coverage for each of the 56 species (termed “strict data set”) had a superalignment length of 170,140 amino acid positions and included 446 gene partitions. Details on data matrix diagnostics are provided in the Supplementary Text, Additional File 1, Supplementary Table S10, Additional File 2, and Supplementary Figures S3-S7, Additional File 3.

### Phylogenetic relationships of sea slug taxa

All analyses irrespective of the data set indicate maximum statistical support for all major splits and phylogenetic relationships on family level (Fig. 1, Supplementary Figures S8-S12, Additional File 3). Notably, low statistical support was inferred with regard to the phylogenetic position of the genus *Embletonia*. In the following, we discuss taxa relationships using the names according to the latest changes [50] that are implemented in World Register of Marine Species [51, 52], although we disagree with several assignments as discussed below.

**Figure 1:**
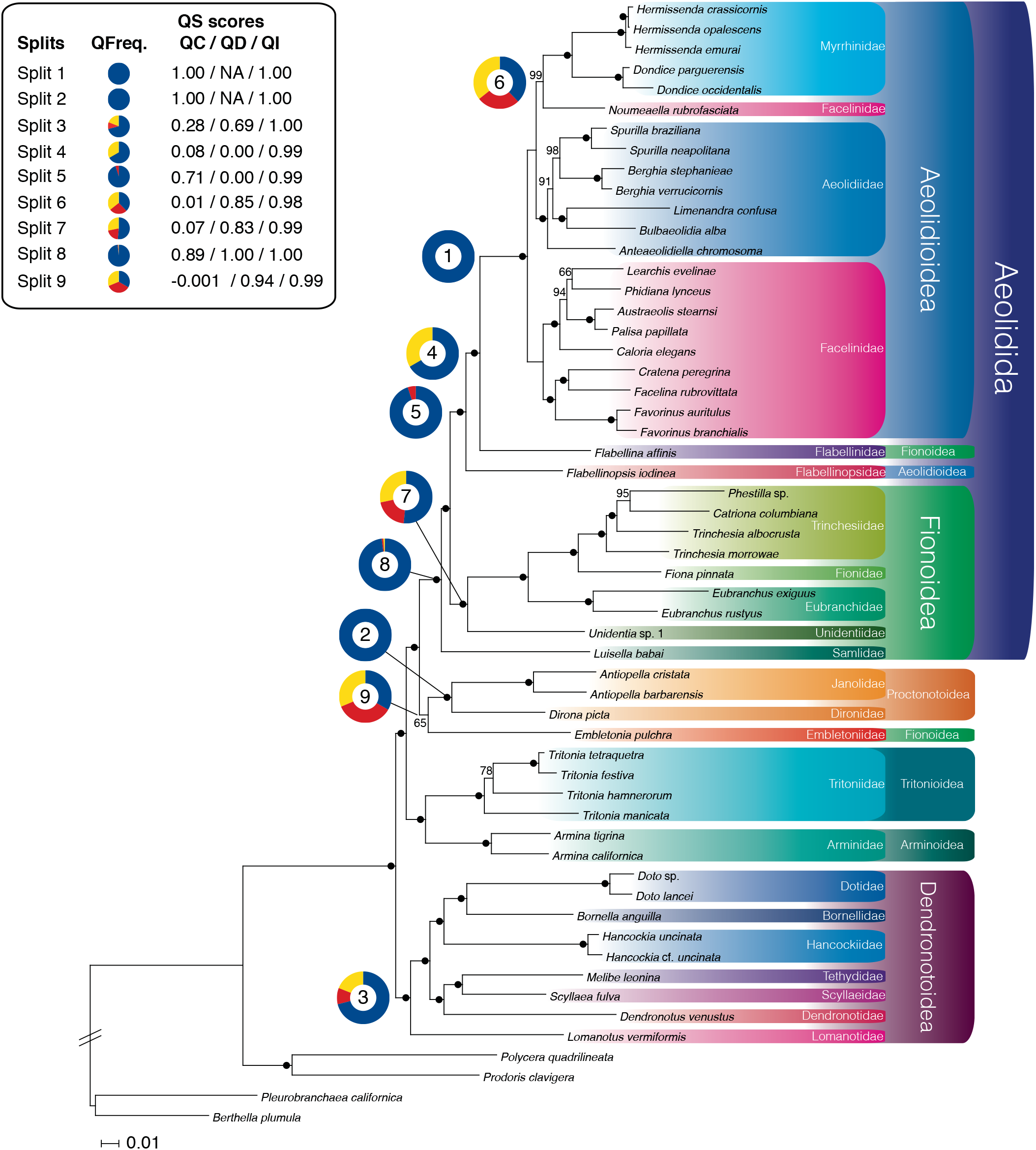
Best ML tree (phylogram) from the strict data set. Maximum likelihood (ML) tree with bootstrap (BS) support values calculated on the strict data set. Black dots (•) indicate a BS support value of 100. The numbers represent splits that are discussed in the main text and the surrounding coloured circles represent Quartet Sampling (QS) scores for the corresponding split. QFreq. = Quartet frequencies. QC = Quartet concordance. QD = Quartet differential. QI = Quartet informativeness.

#### Phylogenetic relationships of major taxa and sea slug families

Out of the seven accepted superfamilies of Cladobranchia, we were able to include members of six superfamilies, whereas a representative of the rare Doridoxoidea was not available to us. We inferred Aeolidida, Aeolidioidea (sensu WoRMS), Proctonotoidea, and Dendronotoidea, with representatives of various families and genera, as being monophyletic. This was fully supported by the quartet scores [41] for Aeolidida, Aeolidioidea, and Proctonotoidea, and strongly supported for Dendronotoidea (see QuartetSampling scores, splits 1-3 and 8 in Fig. 1 and Supplementary Table S11, Additional File 2). Arminoidea and Tritonioidea are only represented by one genus each. Therefore, their assumed monophyly still has to be tested by including relevant genera like *Doridomorpha* in Arminoidea, or *Tochuina* in Tritonioidea.

Our analyses revealed the following ambiguities: *Flabellina affinis* (Flabellinidae), which is currently regarded as a representative of Fionoidea [18], is inferred as sister taxon to Aeolidioidea with maximal statistical support. Quartet sampling, on the other hand, showed only medium support (split 4 in Fig. 1, Supplementary Table S11, Additional File 2) with the large majority of quartets (67%) supporting the focal branch (Aeolidioidea + *Flabellina affinis*), but the strong skew in discordance (quartet differential (QD) = 0) indicating the possibility of a single different evolutionary history supported by all remaining quartets.

The family Flabellinopsidae is currently listed as a member of the Aeolidioidea in WoRMS [52] with *Flabellinopsis iodinea* (Flabellinopsidae) being sister to all remaining taxa in this large clade, confirming previous results [18, 26–28]. Again, this position is statistically maximally supported by classic support values in our study and quartet puzzling scores confirmed this position (split 5, Fig. 1) with strong support (94% of the non-uncertain quartets). Although a strong skew in discordance (QD = 0) indicates the possible presence of an alternative quartet relationship, this result is rather less meaningful due to the low number of discordant trees (5% of the non-uncertain quartets). Thus, our results on Flabellinidae and Flabellinopsidae partly contradict recent analyses and subsequent systematic assignments.

Within Aeolidioidea, the families Myrrhinidae and Aeolidiidae form a monophyletic sister group relationship in our study, thus confirming the results of [28] and [18]. This is also consistent with recent morphological and molecular analyses [53].

The majority of the family Facelinidae is inferred as being monophyletic, but the facelinid species *Noumeaella rubrofasciata* groups with Myrrhinidae in published analyses [18, 28] as well as in our study with nearly maximal ‘classical’ statistical support. However, quartet puzzling only shows weak support for this relationship (38% of the non-uncertain quartets; see split 6 in Fig. 1 and Supplementary Table S11, Additional File 2). In fact, the quartet frequencies show no clear signal since all three quartet topologies are roughly equally supported (27% of the non-uncertain quartets support the second possible quartet topology, 36% support the third; QD = 0.85). Thus, the assignment of this species to Facelinidae [50] or Myrrhinidae (our results) should be reconsidered in future studies. Interestingly, this species did not cluster with other *Noumeaella* species in a three-gene analysis of Aeolidida by Schillo and colleagues [37].

Fionoidea in the sense of Bouchet and colleagues [50] is paraphyletic, mainly due to the position of *Flabellina affinis* and *Embletonia*, the latter is discussed below.

Within Fionoidea, the family Trinchesiidae represented here with three genera, is monophyletic. Unidentiidae is sister to all remaining taxa within Fionoidea. Previously, Korshunova and colleagues [26] inferred this family as sister taxon to Facelinidae and Aeolidiidae. Quartet puzzling analyses, however, do not unambiguously support the relationship of the Unidentiidae as sister to all other Fionoidea. There is rather weak support (52% of the non-uncertain quartets) for said topology and the support for the other two possible quartet topologies is almost similar (QD = 0.99), which indicates that no alternative history is favoured (see split 7 in Fig. 1 and Supplementary Table S11, Additional File 2). In this context, the results of Goodheart and colleagues [18] are quite noteworthy, because in their study, Unidentiidae is the sister taxon of *Embletonia* and the clade *Embletonia* + Unidentiidae is sister to all remaining Fionoidea. Results by Martynov and colleagues [53] suggest a sister group relationship to other aeolidacean families, which is incompatible with our results (see below).

The family Samlidae, represented by *Luisella babai*, is considered as being part of Fionoidea [18]. In our study, however, it is inferred as sister to all remaining Aeolidida in all analyses with maximum ‘classical’ tatistical support as well as very strong quartet support (see split 8 in Fig. 1 and Supplementary Table S11, Additional File 2): About 98% of the quartets supported this relationship, without evidence for alternative quartet topologies (QD = 1), confirming previous results by Korshunova and colleagues [26].

With regard to Proctonotoidea, Tritonioidea, and Dendronotoidea, our results confirm the findings published by Goodheart and colleagues [18] with the family Embletoniidae being the only exception, as we will discuss below.

### The phylogenetic position of Embletoniidae remains ambiguous

The monogeneric family Embletoniidae, which currently only comprises two recognized species, *Embletonia pulchra* and *E. gracilis*, has experienced a vivid history since the first description of the genus *Embletonia* by Alder and Hancock [54], with *Pterochilus pulcher* Alder and Hancock, 1844 as type species. The authors considered this species as a link between cladobranch aeolids and panpulmonate sacoglossans, two taxa that are not closely related to each other, but show many convergent characters. Pruvot-Fol [31], who named the family for the first time, included members of Trinchesiidae, but assigned the whole clade as a “section” to the dendronotoid family Dotidae. The two recognized members of *Embletonia* share some characters with members of Fionoidea or Aeolidioidea, e.g., the reduction of the lateral teeth, the absence of rhinophoral sheaths [56], and the presence of a cnidosac at the end of the cerata, a synapomorphy of Aeolidida [19], which additionally favours a position within this clade. However, Martin and colleagues [16] and Goodheart and colleagues [18] have shown that this cnidosac differs to a great extent from the typical aeolidid cnidosac by lacking a proper sac-like structure with musculature around it, as well as a connection to the digestive gland, which is necessary for taking up sequestered cnidocysts. Nevertheless, cnidocysts were found in the structures investigated by Goodheart and colleagues [18]. The authors explain this atypical situation with a loss of characters or as constituting a transitional form in the evolution of the cnidosac. Most recently, Martynov and colleagues [53] provided evidence for paedomorphic processes, which would explain a regressive evolution within Embletoniidae. This phenomenon is quite common in various unrelated taxa inhabiting soft-bottom interstitial environments. *Embletonia* feeds on hydrozoans, which is a typical food source of many aeolidids, but also of some dendronotoids. Unique to this genus are the cerata, which show bi- to quadrifid apices. Highly structured cerata are not known from any aeolidids. However, the digestive gland reaches far into these cerata, a character less pronounced in Proctonotoidea, and only present in one further non-aeolidid group, the genus *Hancockia*.

*Embletonia* also shares traits that are characteristic for non-aeolidid groups, a reason why Pruvot-Fol [31] included the genus into the family Dotidae (Dendronotoidea). This assignment to Dotidae, as well as grouping with Trinchesiidae was, however, rejected later by Schmekel [32], and the closer relationship to Dendronotoidea was emphasized by Miller and Willan [57]. The primary connecting character is the lack of oral tentacles, which are considered to be a synapomorphy of the Aeolidida [19]. Furthermore, their oral gland ducts do not open into the oral tube by two separate ducts, but fuse into one common duct, which is described for Proctonotoidea. Proctonotoidea mainly feed on bryozoans, however, a few members also rely on hydrozoan prey, similar to *Embletonia*.

Few studies addressed the phylogenetic relationship of Embletoniidae using molecular data [18, 24, 53]. Bleidissel [24] focussed on Aeolidida and included *Embletonia*, because of its putative assignment to this group. Bleidissel’s analyses, based on three genes, inferred a sister group relationship of Embletoniidae with Notaeolidiidae, with the latter again being sister to all remaining Aeolidida. In the only study based on a large data set, *Embletonia* was inferred, along with *Unidentia*, within Aeolidida as sister to the remaining Fionoidea, thus excluding a closer relationship with *Notaeolidia* [18]. Martin and colleagues [16] included characters of the cnidosac into the morphological data matrix published by Wägele and Willan [19], and their analysis resulted in an assignment of *Embletonia* to Aeolidida (tree not shown in the publication). Likewise, our unpublished morphological analyses render *Embletonia* as a sister taxon to Aeolidida. However, it is more likely the lack of data that constrains the position than apomorphic characters of high phylogenetic information.

In our analyses comprising the unreduced and strict data set, *Embletonia pulchra* is inferred as sister to Proctonotoidea, but with negligible support in the strict data set (65 BS, 50.1 aLRT, 1 aBayes). When assuming that *Embletonia* is a sister taxon of the Proctonotoidea (see split 9 in Fig. 1 and position i in Fig. 2) and taking into consideration the studies on the evolution of prey preferences [28] and cnidocyst incorporation [18], we have to conclude that (1) feeding on Hydrozoa is an old trait within Cladobranchia and has not changed in *Embletonia* (in contrast to Proctonotoidea) and (2) the evolution of the cnidosac might have started in the stemline of the clade Aeolidida/Proctonotoidea/Embletoniidae, with *Janolus* and *Dirona* probably representing a condition where the ability to store cnidocysts was lost due to a food switch to bryozoan prey. Both, an independent evolution of cnidosacs and cnidocyst storage (in the genus *Hancockia*) as well as a loss or strong reduction of these complex structures has occurred within Dendronotoidea [18].

**Figure 2:**
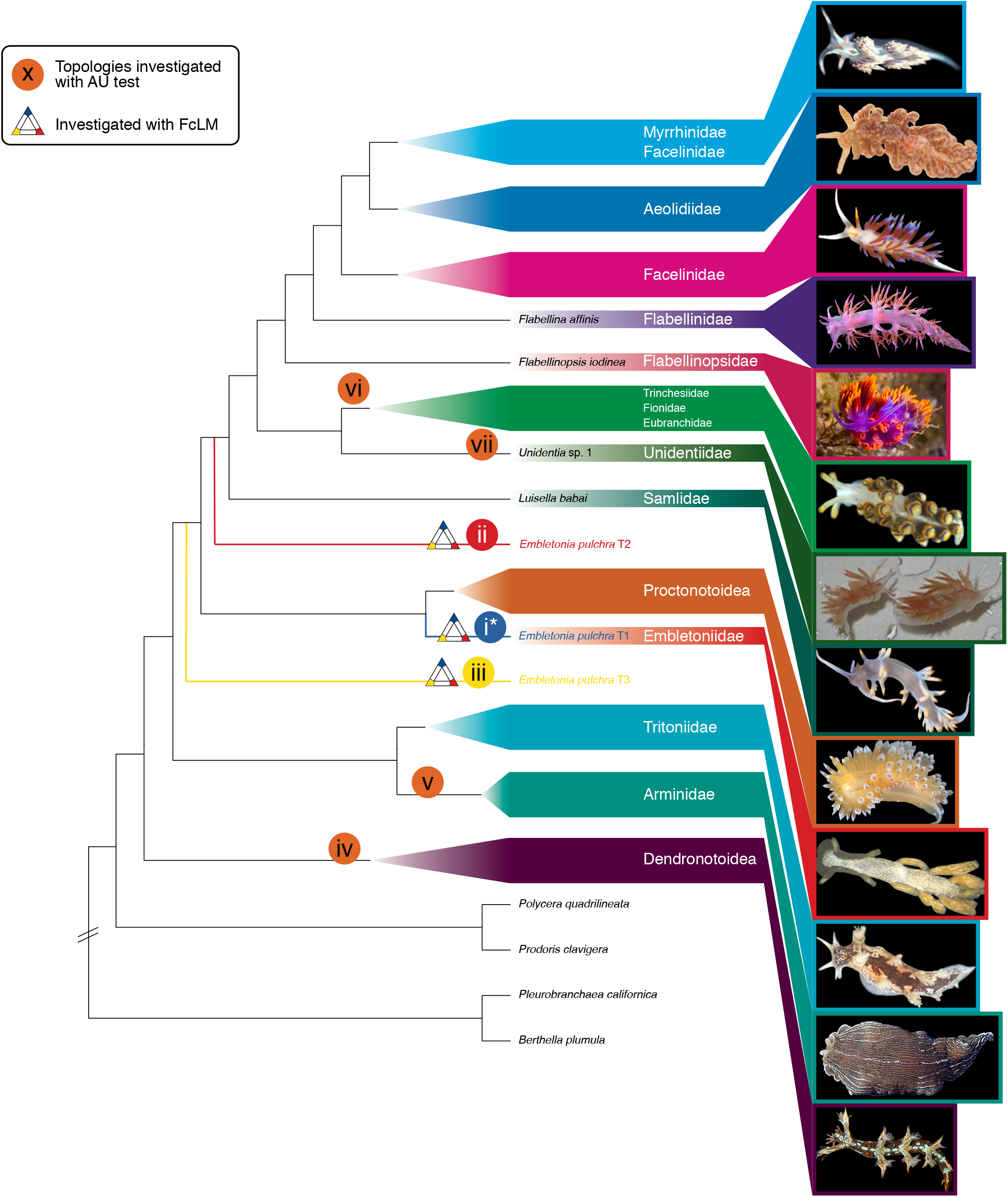
Best ML tree (cladogram): AU tests + FcLM. Cladogram with summarized major families/clades and images of representative species. Splits for which additional testing was performed are marked with Roman numerals (i-vii) in a coloured circle (AU test) and a triangle (FcLM, splits i-iii). The original position of *E. pulchra* as obtained from the strict data set is marked by a blue branch (T1). Alternative positions of *E. pulchra* are indicated by a red (T2) and yellow branch (T3), respectively. We thank Craig A. Hoover for providing the picture of *Flabellinopsis iodinea* and Karen Cheney for permissions to use the picture of *Unidentia angelvaldesi*.

In our results from the intermediate data set, *Embletonia* is a sister group to all remaining Aeolidida, but with even less support (51 BS, 33.1 aLRT, 1 aBayes). Considering this relationship as a possible evolutionary scenario (Fig. 2, position ii, results on the intermediate data set) means that the evolution of the cnidosac would have had to start in the stemline of Embletoniidae/Aeolidida, while the typical character of the Dendronotoidea, the rhinophoral sheaths, had already been lost and oral tentacles had not yet evolved.

However, both discussed possibilities (see Fig. 2, positions i and ii) are neither supported statistically by classical bootstrap values, nor by our quartet analyses: Frequencies of the three possible quartet topologies are almost equal (33% vs. 35% vs. 31% of all non-uncertain quartets, split 9 in Fig. 1 and Supplementary Table S11, Additional File 2), which indicates a highly complex evolution or rapid radiation.

Morphological analyses of important characters, like the positions of the anus, jaws, and radula also contradict both relationships discussed above with apomorphic features lacking for both hypotheses [53]. Instead, Embletoniidae shows an uniserial radula with central teeth more similar to various aeolidids.

### Evaluation of alternative positions of Embletoniidae and possible confounding signal

To gain more insights into one of the obtained positions of *Embletonia* and to investigate alternative positions (see Fig. 2), further analyses were conducted. Note, that we consider the strict data set as most reliable, since it has full gene coverage for all species, following the rationale of Dell’Ampio and colleagues [58] and Misof and colleagues [40], who showed that inferred positions with high statistical support can be simply due to non-phylogenetic signal, e.g., the distribution of missing data. However, we also performed some of the analyses on the intermediate data set.

We applied approximately unbiased (AU) tests [38] for alternative positions of *Embletonia* on the intermediate and strict data set. An AU test always takes the complete tree topology into account and not only single splits. Further, it does not test whether or not confounding signal is inherent in the data set, e.g., due to non-randomly distributed data and/or among-lineage heterogeneity violating SRH conditions. We therefore also applied Four-cluster Likelihood Mapping (FcLM) [39] along with a permutation approach on the strict data set. By testing all three possible quartet topologies around *Embletonia* we evaluated whether or not there was an alternative signal. Further, we checked for any sign of confounding signal (see [40]). To this end, we defined four groups (Supplementary Table S12, Additional File 2) considering group 4 as outgroup. We performed separate analyses for two outgroup variations: first, with 19 species including Anthobranchia and Pleurobranchomorpha and second, only with the 15 remaining cladobranch species. We drew quartets on the original data set and on three artificial data sets, from which any existing phylogenetic signal was removed in three different ways (see Materials and methods, Supplementary Text, Additional File 1, and [40]): (a) by destroying the phylogenetic signal but leaving the distribution of missing data and the compositional heterogeneity, which can lead to violating SRH conditions, untouched; (b) by leaving the distribution of missing data untouched but making the data set completely homogenous (no SRH model violation possible), and (c) by randomizing the missing data distribution and making the data set completely homogenous. For all details see Supplementary Text, Additional File 1.

Interestingly, the results of the phylogenetic trees and the results of the FcLM (Supplementary Table S13, Additional File 2) and AU tests (Supplementary Table S14, Additional File 2) were quite contradicting:

i. Although the ML trees of the unreduced and strict data sets suggest that *Embletonia* is sister to Proctonotoidea and although the AU test was unable to reject this topology (p > 0.05), it received the lowest proportion of quartets (< 20%) in the FcLM approach. Thus, this relationship can only be explained by confounding signal (see original and permutation results in Supplementary Table S13, Additional File 2).
ii. Although the best ML tree of the intermediate data set suggests *Embletonia* to be sister to all remaining Aeolidida, a position that is not rejected by the AU test (p > 0.05), the FcLM results indicate only minimal support for such a relationship: the proportion of supporting quartets, excluding those that can be explained by confounding signal, was only around 3%. This also implies that AU tests, irrespective of whether or not a topology for the data set is significantly rejected, cannot be used to check if the signal is confounding.
iii. A sister group relationship of *Embletonia* to a clade Aeolidida + Proctonotoidea, which received strongest support in the FcLM analyses (8-16% of all quartets after excluding the proportion of supporting quartets that can be explained by confounding signal, see Supplementary Table S13, Additional File 2), was equally rejected by the AU test. There is only very little signal that is not confounding (around 3-8%, compare quartets of original with permuted approaches, Supplementary Table S13, Additional File 2), which would support either *Embletonia* + Aeolidida (position ii in Fig. 2) or *Embletonia* as sister to a clade Aeolidida + Proctonotoidea (position iii in Fig. 2). Thus, these results clearly indicate that the position of *Embletonia* as a sister taxon of Proctonotoidea is not due to any informative phylogenetic signal, but only due to confounding signal in our data set, and again leaves the phylogenetic position of *Embletonia* as an enigma. In order to analyse further possibilities of putative relationships of *Embletonia*, we tested four alternative positions (iv - vii, see Fig. 2) of *Embletonia*, which have been discussed in the literature before, by applying the AU test on the strict data set (see Fig. 2 and see below). Note that none of these positions were inferred in any of our ML analyses.
iv. Since *Embletonia* exhibits characters, which are shared with the Dendronotoidea, we analysed a putative sister group relationship with this superfamily.
v. Although an assignment to Tritonioidea is very unlikely, because *Embletonia* does not share all the characters special for this superfamily, the position of the Arminoidea is variable within the various published phylogenies [18, 59, 60] when including this superfamily. Nevertheless, we tested this possibility.

The last two tests imply a closer relationship of *Embletonia* with Fionoidea, a relationship that was assumed in former times and reflects the current systematics [50]. Therefore, we tested (vi) a position of *Embletonia* as sister to Fionoidea and (vii) *Embletonia* as sister to Unidentiidae and this clade being again sister to the remaining Fionoidea in restricted sense [18, 53].

AU tests significantly rejected (p < 0.05) all four alternative positions (iv - vii, see Fig. 2) of *Embletonia* (see Supplementary Table S14, Additional File 2).

Despite our extensive molecular data sets and tests, we still cannot unambiguously assign *Embletonia* to one of the superfamilies in our tree. Beyond only small putative phylogenetic signal as indicated by our FcLM analyses, which is also in line with the negligible support considering classical statistical support, a reason could be the lack of relevant taxa in our data set that could positively influence the position of Embletoniidae in the cladobranch tree (e.g., Doridomorpha, Charcotiidae, Notaeolidiidae). Interestingly, morphological traits are also confounding and do not yet allow for an unambiguous assignment. Because of its unresolved position, several evolutionary traits within the Cladobranchia cannot be satisfactorily explained.

## Conclusions

Due to the high number of orthologous single-copy genes that could be successfully extracted from the transcriptomes, the high information content and up to full gene coverage of the supermatrices, and the high resolution of all three phylogenies, we conclude that the use of transcriptomic data is a valuable tool for analysing phylogenetic relationships within Cladobranchia. Nevertheless, analyses of large data sets can be error-prone to systematic bias and classical support values might be inflated as has been shown and discussed [61–64]. Beyond careful data processing prior to phylogenetic tree inference, additional thorough tests, e.g., AU tests, quartet approaches like FcLM and quartet puzzling as well as checks for confounding signals on a variety of different data matrices become more and more indispensable. Our study has revealed that, despite previous efforts, the position of some families within this group, especially the Embletoniidae, requires further investigation and possibly taxonomic revision. In future studies, the present data set should be extended by increasing the number of group-specific orthologous single-copy genes and by including

Charcotiidae, Notaolidiidae and other relevant species to shed light on the relationships between families and superfamilies in Cladobranchia in order to draw a more complete image of the evolution of this enigmatic group.

## Materials and methods

An overview of the complete workflow is displayed in Fig. 3. Major steps are described here while all details and settings can be found in the Supplementary Text, Additional File 1.

**Figure 3:**
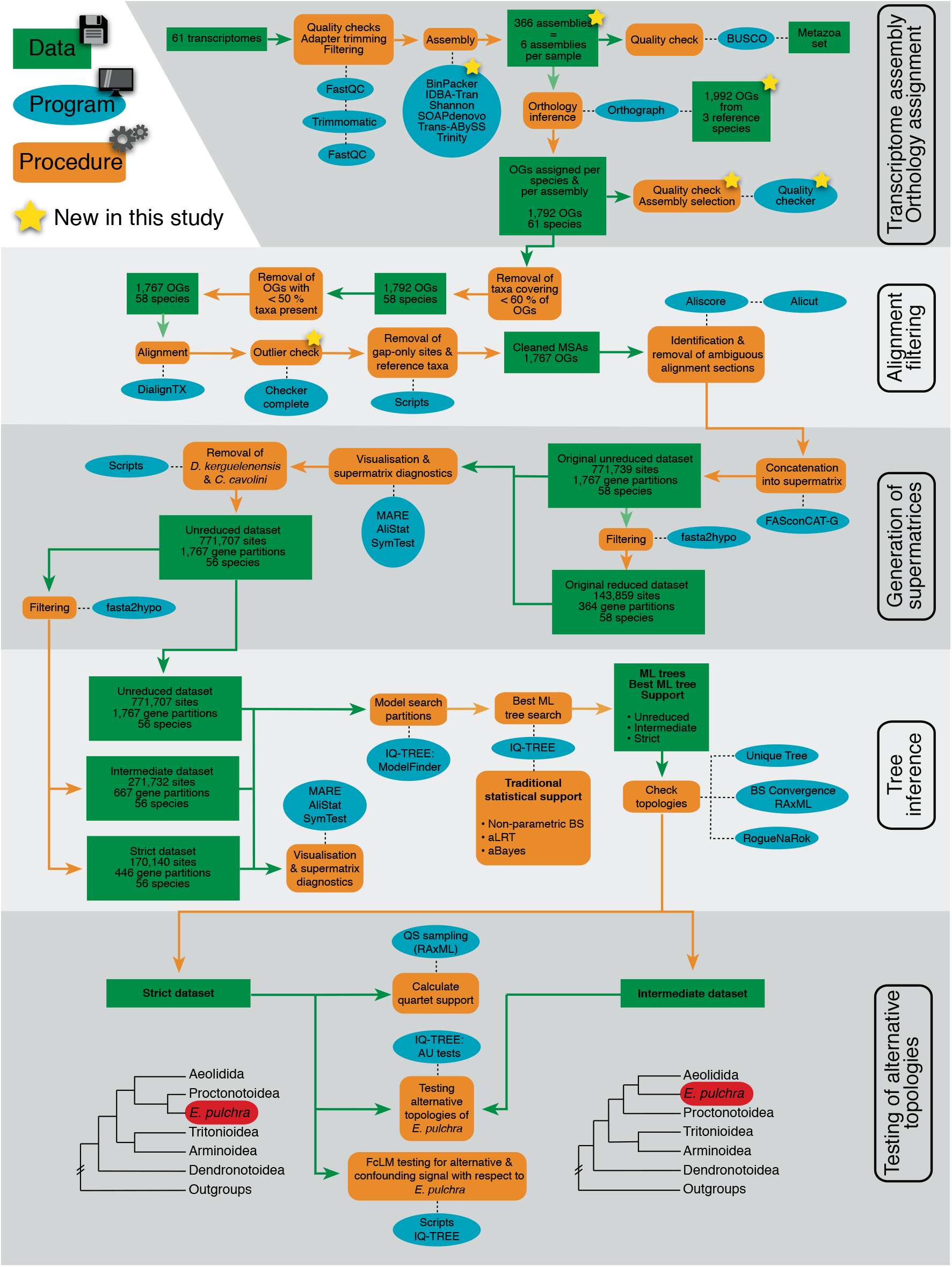
Analysis workflow. Schematic workflow representing all steps from NGS data to the testing of alternative topologies with major steps being highlighted in shades of gray.

### Taxon sampling and sampling of transcriptome data

For this study, we used recently published transcriptome data and generated new transcriptome data for 21 species. We collected 19 species of Cladobranchia and two more distantly related species of heterobranch sea slugs from different locations in the Mediterranean Sea and the Sea of Japan (Supplementary Table S1, Additional File 2). The specimens were preserved in RNAlater (Qiagen) or IntactRNA (Evrogen) and stored at −80 °C. The specimens collected on Elba island (Supplementary Table S1, Additional File 2) were stored at −20 °C for approximately two weeks and then transferred to −80 °C until RNA extraction. RNA extraction was performed using the Macherey & Nagel NucleoSpin RNA II kit. Preparation and amplification of the cDNA libraries were performed by StarSeq GmbH, Mainz using the Illumina TruSeq Stranded RNA HT kit. Paired-end sequencing was also conducted at StarSeq with a read length of 150 base pairs on an Illumina NextSeq 500 sequencing platform. Raw reads were submitted to the NCBI SRA database. All accession numbers are provided in Supplementary Table S2, Additional File 2.

Our newly generated transcriptome samples were combined with the published transcriptome data of another 40 samples that we downloaded from the NCBI SRA database (Supplementary Table S2, Additional File 2) [27, 28, 65, 66]. The published data comprised 37 species of Cladobranchia as well as two dorids, *Prodoris clavigera* and *Doris kerguelenensis*, and one pleurobranchid, *Pleurobranchaea californica* (Supplementary Table S2, Additional File 2).

### De novo transcriptome assembly

All raw sequence reads of published and newly generated samples were quality-checked prior to and after adapter trimming using FastQC Version 0.11.5 [67]. Adapter trimming and quality filtering were performed with Trimmomatic v0.36 [68] using a custom adapter file (see Additional File 4).

Data from altogether 61 samples were assembled using six assembly tools: BinPacker v. 1.1 [69], IDBA-Tran v. 1.1.1 [70], Shannon v 0.0.2 [71], SOAPdenovo-Trans v. 1.04 [72], Trans-ABySS v. 1.5.5 [73], and Trinity v. 2.4.0 [74]. All assemblers were run with default settings and all paired-end reads that survived the trimming process were used as input. We additionally provided surviving single-end reads to those assemblers that were capable of processing them (IDBA-Tran, SOAPdenovo-Trans, and Trans-ABySS).

Following identification of the best transcriptome assembly per species (see below), possible foreign contaminants were identified upon submission of the newly sequenced transcriptomes to NCBI Transcriptome Shotgun Assembly (TSA) database and subsequently removed from the sequences. Details are provided in the Supplementary Text, Additional File 1 and in Supplementary Table S7, Additional File 2. The five alternative assemblies for each sample that has been sequenced in frame of this study are provided in Additional File 5.

### Orthology prediction and generation of data matrices

We designed a custom-made orthologue set by selecting all genes that were listed by OrthoDB version 9 [75] to be single-copy at the hierarchical level “Lophotrochozoa” and downloaded the respective table with the IDs of the orthologue groups (called OGs hereinafter). We additionally downloaded the official gene sets of three species with well-sequenced and annotated genomes, which we selected as reference species (i.e. *Biomphalaria glabrata*, Official Gene Set (OGS) version 1.2 vectorbase [76], *Crassostrea gigas*, OGS version Sep-2012 (ENA genebuild) [77], and *Lottia gigantea*, OGS version Jan-2013 (JGI genebuild) [78]. We excluded five genes from this set due to defective sequence headers, leading to a custom-made orthologue set comprising 1,992 orthologues. Orthology prediction was performed using Orthograph v.0.6.2 [79], for which we used the aforementioned orthologue set (Additional File 6). Details are provided in the Supplementary Text, Additional File 1. To reduce the amount of missing data per species, three transcriptome assemblies that covered less than 60% of the orthologue set were excluded from further analyses: *Pseudobornella orientalis* (53% of the orthologue set missing), *Dermatobranchus* sp. (46% missing), and *Tritoniopsis frydis* (51% missing). We then removed all OGs for which less than 50% of the investigated species had a positive hit. This resulted in 1,767 OGs for further analyses.

The quality of all transcriptome assemblies was further assessed with BUSCO v3.0.0 using the metazoa_odb9 reference set genes comprising 978 BUSCO groups [42] and default settings (Supplementary Table S5, Additional File 2). Because BUSCO’s general metazoa data set is not very specific for nudibranchs and since there is no way to easily compile a nudibranch-specific reference data set (R. Waterhouse, personal communication), we devised a method that makes use of the output generated by Orthograph. For each Orthograph run, we calculated the number of sequences that were assigned to OGs by Orthograph as well as the cumulative length of these sequences. With the aim to maximize the amount of data, the latter was used as a criterion to determine the best assembly for each species (for details see Supplement Text, Additional File 1, Supplementary Table S6, Additional File 2, and Additional File 7).

Multiple sequence alignments on translational level were generated using DIALIGN-TX Version 1.0.2 [80] and checked for outlier sequences using a newly implemented version of the outlier script described in [40] (see Supplementary Text, Additional File 1 for details; unfiltered alignments are provided in Additional File 8). Sequences identified as outliers as well as all sequences belonging to the three reference taxa were removed from the alignments (Additional File 9).

The amino acid multiple sequence alignments were examined with the program Aliscore version 2.0 [81, 82] in order to identify ambiguous or randomly similar aligned sites. All positions flagged by Aliscore (~ 29% of the originally aligned sites, see Supplementary Text, Additional File 1) were discarded using AliCut version 2.31 [83] (Additional File 10). The resulting masked amino acid alignments were concatenated into a supermatrix along with the creation of a partition file using FASconCAT-G version 1.04 [84].

### Compilation, evaluation and optimization of data sets

This amino acid supermatrix, with 58 species and including 1,767 genes, was analysed using the software tool MARE version 1.2-rc [44] in order to assess the potential information content (IC) of each gene partition, the overall information content of the matrix, and the coverage in terms of gene partitions. The tool AliStat version 1.6 [45] was used to calculate alignment diagnostics and the software SymTest version 2.0.47 [46–48] was used to analyse the compositional heterogeneity of the supermatrix in order to detect possible violations of stationary, (time-)reversible, and homogeneous (SRH) conditions [49].

To reduce especially among-lineage heterogeneity (see Results and Discussion), we excluded the species *Doris kerguelenensis* and *Calmella cavolini* from our data (see Supplementary Figures S2 and S7, Additional File 3).

We repeated analyses with MARE, AliStat, and SymTest and compiled three final data sets, allowing different levels of missing data (Supplementary Table S10, Additional File 2): an unreduced data set with 56 species and all 1,767 gene partitions with 771,707 aligned amino-acid sites and allowing ~ 39% missing data; an intermediate data set in which data for at least one representative of the defined groups (Supplementary Table S9, Additional File 2) had to be present, which led to a data matrix of 56 species and 667 gene partitions (271,732 aligned sites) with 98% gene coverage and 18% of missing data, and our most strict data set only including genes present in all 56 species. This led to a data matrix with 170,140 aligned sites, 446 gene partitions and less than 13% of missing data. Missing data can lead to confounding signals in phylogenetic inference [40, 44, 58]. We therefore consider our strict data set as most reliable. Details are provided in the Supplementary Text, Additional File 1. The three supermatrices are provided in Additional File 11.

### Phylogenetic tree inference

For all three data sets, maximum likelihood (ML) trees were calculated using IQ-TREE version 1.6.12 [85]. The best fitting amino acid models for each partition were identified using ModelFinder [86], which was run using an edge-link partitioned approach [87]. Out of 20 tree searches per data set, we selected the best ML tree according to the best loglikelihood. Statistical support was derived from non-parametric bootstrap replicates ensuring bootstrap convergence. Additionally, we calculated SH-like approximate likelihood ratio test support [88] and approximate Bayes test support [89]. The best ML tree of each of the three data sets was tested for the presence of rogue taxa using RogueNaRok v.1.0 [90]. Details for each step including used settings are provided in the Supplementary Text, Additional File 1.

### Testing for alternative topologies

#### Quartet puzzling

To analyse phylogenetic discordance, we applied the Quartet Sampling (QS) method [41], which aims to identify the lack of branch support due to low phylogenetic information, discordance due to lineage sorting or introgression, and misplaced or erroneous taxa (rogue taxa). Details on the analysis and interpretation of scores are provided in the Supplementary Text, Additional File 1 and Supplementary Table S11, Additional File 2.

#### Testing the position of Embletonia

Since the inferred position of *Embletonia pulchra* was not stable comparing the best ML trees of the intermediate and strict data set, we tested various possible topologies with AU tests (see Fig. 2) [38] as implemented in IQ-TREE version 1.6.12 (see Results and Discussion, Supplementary Text, Additional File 1, and Additional File 12). To further analyse whether or not the placement of *Embletonia* in our best tree inferred from the strict data set was influenced by confounding signal and violating SRH conditions, and whether or not there was putative phylogenetic signal for alternative positions of *Embletonia*, we additionally performed Four-cluster Likelihood Mapping (FcLM), which is outlined in the results section and in detail in the Supplement Text, Supplementary File 1 (see also Additional File 13). In summary, we tested the following seven alternative hypotheses concerning the position of *Embletonia*:

i. *Embletonia* is sister to Proctonotoidea (AU test + FcLM)
ii. *Embletonia* is sister to all Aeolidida (AU test + FcLM)
iii. *Embletonia* is sister to (Aeolidida, Proctonotoidea) (AU test + FcLM)
iv. *Embletonia* is sister to Dendronotoidea (AU test)
v. *Embletonia* is sister to Arminoidea (AU test)
vi. *Embletonia* is sister to Fionoidea (AU test)
vii. *Embletonia* is sister to Unidentiidae and this clade is sister to remaining Fionoidea (AU test).

## Abbreviations

aLRT: approximate likelihood ratio test
AU test: approximately unbiased test
BS: bootstrap
FcLM: Four-cluster Likelihood Mapping
IC: information content
ML: maximum likelihood
MSA: multiple sequence alignment
OG: orthologue group
OGS: official gene sets
QD: Quartet differential
QS: Quartet Sampling
SRH: stationary, (time-)reversible and homogenous
TSA database: Transcriptome Shotgun Assembly database, WoRMS:

## Declarations

### Ethics approval and consent to participate

Not applicable

### Consent for publication

Not applicable

### Availability of data and materials

The data sets and scripts supporting the conclusions of this article are available via Figshare, “[UNIQUE PERSISTENT IDENTIFIER AND WEB LINK TO DATA SET(S) WILL BE PROVIDED UPON ACCEPTANCE OF THE ARTICLE].”

### Competing interests

The authors declare no competing interests.

### Funding

AD and HW received funding by the German Research Foundation (DFG DO 1781/1-1, DFG WA 618/18-1). AM received support by the MSU Zoological Museum (AAAA-A16-116021660077-3). The work of TK was conducted under the IDB RAS Government basic research program in 2020 No. 0108-2019-0002.

### Authors’ contributions

AD, DK, and HW designed the study. DK, HW, MS, AM, and TK collected and provided material. DK, KM, AD, and HW performed all data analyses. DK and AD developed scripts. AD performed sequence data management. MS, AM, and TK provided pictures. JG provided access to unpublished data. All authors contributed in writing the manuscript, with AD, DK, KM, and HW taking the lead. All authors read and approved the final manuscript.

## Acknowledgements

KM thanks Ondrej Hlinka (CSIRO) for HPC support and Thomas Wong (ANU) for kindly providing Unique Tree. KM and DK thank Robert Waterhouse for details on official gene sets used in OrthoDB 8. DK thanks Malte Petersen for suggestions on using Orthograph.

## Additional Files

### Additional file 1 - Supplementary Text (pdf)

### Additional file 2 - Supplementary Tables S1 - S14 (xlsx)

**Table S1:** Sampling information for the species collected for this study.

**Table S2:** NCBI accession numbers for all species used in this study.

**Table S3:** Statistics of raw sequence reads before and after trimming.

**Table S4:** Assembly statistics.

**Table S5:** BUSCO results.

**Table S6:** Results of the Quality Checker script and selection of the best assembly.

**Table S7:** Information on sequences removed during contamination filtering.

**Table S8:** Number of removed outlier sequences per species.

**Table S9:** Group definitions to compile the intermediate data set.

**Table S10:** Supermatrix diagnostics of data sets used in this study.

**Table S11:** Results of the Quartet Sampling analysis.

**Table S12:** Group definitions used for Four-cluster Likelihood Mapping (FcLM) analyses.

**Table S13:** FcLM results testing the position of *Embletonia*.

**Table S14:** AU test results on the strict and intermediate data set.

### Additional file 3 - Supplementary Figures S1 - S12 (pdf)

**Figure S1: Species-pairwise site-coverage of the original unreduced and reduced data sets.**

Heat maps indicate species-pairwise amino acid site-coverage of the sequences of 58 species in the original data sets inferred with AliStat. Low shared site-coverage is in shades of red and high shared site-coverage is in shades of green. AliStat scores are given in Supplementary Table S10, Additional File 2. **a)** original unreduced data set. **b)** original reduced data set.

**Figure S2: Heat maps calculated with SymTest applying the Bowker’s test on the original unreduced and reduced data sets.**

Heat maps show the results of pairwise Bowker’s test as implemented in SymTest 2.0.47 analysing the original data sets unreduced and reduced. The percentage of pairwise p-values < 0.05 rejecting SRH conditions are given in parentheses. **a)** original unreduced data set (p-values < 0.05: 83.36%). **b)** original reduced data set (p-values < 0.05: 42.65%). Note that especially *Calmella* and *Doris* are obvious with respect to violating SRH conditions.

**Figure S3: Heat map visualising the information content of the final unreduced data set calculated with MARE.**

The information content (IC) is colour-coded in shades of blue, with darker shades representing higher IC and white squares indicating missing data. Red squares indicate gene partitions with an IC = 0. Species are displayed in rows (x-axis) and gene partitions are displayed in columns (y-axis). Supermatrix diagnostics of MARE are provided in Supplementary Table S10, Additional File 2.

**Figure S4: Heat map visualising the information content of the final intermediate data set calculated with MARE.**

The information content (IC) is colour-coded in shades of blue, with darker shades representing higher IC and white squares indicating missing data. Red squares indicate gene partitions with an IC = 0. Species are displayed in rows (x-axis) and gene partitions are displayed in columns (y-axis). Supermatrix diagnostics of MARE are provided in Supplementary Table S10, Additional File 2.

**Figure S5: Heat map visualising the information content of the final strict data set calculated with MARE.**

The information content (IC) is colour-coded in shades of blue, with darker shades representing higher IC and white squares indicating missing data. Red squares indicate gene partitions with an IC = 0. Species are displayed in rows (x-axis) and gene partitions are displayed in columns (y-axis). Supermatrix diagnostics of MARE are provided in Supplementary Table S10, Additional File 2.

**Figure S6: Species-pairwise site-coverage of the final unreduced, intermediate, and strict data set.**

Heat maps indicate species-pairwise amino acid site-coverage of the sequences of 56 species in the final data sets inferred with AliStat. Low shared site-coverage is in shades of red and high shared site-coverage is in shades of green. AliStat scores are given in Supplementary Table S10, Additional File 2. **a)** unreduced data set. **b)** intermediate data set. **c)** strict data set.

**Figure S7: Heat maps calculated with SymTest applying the Bowker’s test on the final unreduced, intermediate, and strict data sets.**

Heat maps show the results of pairwise Bowker’s test as implemented in SymTest 2.0.47 analysing the final data sets unreduced, intermediate, and strict. The percentage of pairwise p-values < 0.05 rejecting SRH conditions are given in parentheses. **a)** unreduced data set (p-values < 0.05: 82.14%). **b)** intermediate data set (p-values < 0.05: 63.96%). **c)** strict data set (p-values < 0.05: 46.17%).

**Figure S8: Best ML tree of the strict data set with aLRT and aBayes support.**

The phylogram is identical to the phylogram in Fig. 1. The first value displays branch support based on 10,000 SH-aLRT replicates, the second value displays support derived from the approximate Bayesian support.

**Figure S9: Best ML tree of the intermediate data set with non-parametric bootstrap support.**

Statistical support was inferred from 300 non-parametric bootstrap replicates.

**Figure S10: Best ML tree of the intermediate data set with aLRT and aBayes support.**

The first value displays branch support based on 10,000 SH-aLRT replicates, the second value displays support derived from the approximate Bayesian support.

**Figure S11: Best ML tree of the unreduced data set with non-parametric bootstrap support.**

Statistical support was inferred from 100 non-parametric bootstrap replicates.

**Figure S12: Best ML tree of the unreduced data set with aLRT and aBayes support.**

The first value displays branch support based on 10,000 SH-aLRT replicates, the second value displays support derived from the approximate Bayesian support.

### Additional file 4 - FASTA file in zip archive

**Archive S1**: Illumina adapters used for adapter trimming.

### Additional file 5 - FASTA files in zip archive

**Archive S2**: Included in this archive are the five alternative assemblies for each sample that has been sequenced in the frame of this study (FASTA format). Note that the best selected assembly has been deposited at the NCBI TSA database. *Pseudobornella orientalis* (HW08) has been removed from the NCBI TSA database due to exceptionally low sequence quality. Its alternative assemblies are therefore also not part of this archive.

### Additional file 6 - FASTA/txt files in zip archive

**Archive S3**: This archive includes official gene sets of the three reference species *Biomphalaria glabrata, Crassostrea gigas*, and *Lottia gigantea* on translational and transcriptional level, the list of all orthologous sequence clusters (OGs) as required for Orthograph, and an exemplary Orthograph config file.

### Additional file 7 – Python script/txt files in zip archive

**Archive S4**: Included in this archive is the *Orthograph_Quality_Checker.py* script, a manual, an example configuration file, and an example output file.

### Additional file 8 – Alignment files in zip archive

**Archive S5:** Unmasked multiple sequence alignments on amino acid level including *Doris kerguelenensis* and *Calmella cavolini* prior to the removal of outliers.

### Additional file 9 – Python scripts in zip archive

**Archive S6**: This archive contains two custom Python scripts. The *remove_outliers.py* script removes all identified outlier sequences from a given alignment. The *remove_reference_sequences.py* script removes all sequences from the reference species *Biomphalaria glabrata, Crassostrea gigas*, and *Lottia gigantea* from the alignments.

### Additional file 10 – Alignment files in zip archive

**Archive S7**: 1,767 Multiple sequence alignments (FASTA format) on amino acid level, from which sequences belonging to *Doris kerguelenensis* and *Calmella cavolini* as well as ambiguously aligned sections and gap-only sites were removed. These served as the basis for compiling the final unreduced supermatrix.

### Additional file 11 – Alignment/txt files in zip archive

**Archive S8**: The unreduced, intermediate, and strict supermatrix (FASTA format) plus respective gene partition information including the selected substitution model used in the phylogenetic analyses.

### Additional file 12 – Tree files (NEWICK format) in zip archive

**Archive S9:** Seven tree topologies (NEWICK format) displaying differing positions of *Embletonia pulchra* that were tested using the approximately unbiased (AU) test with IQ-TREE.

### Additional file 13 - Alignment/NEXUS/txt files in zip archive

**Archive S10:** Data used for Four-cluster Likelihood Mapping (FcLM). This archive includes two directories (one per approach, with a) 19 species included in Group 4 and b) 15 species included in Group 4; see section 17). Each directory includes four subdirectories: original, permutationI, permutationII, and permutationIII. In each subdirectory, the following files that served as input for the FcLM with IQ-TREE are provided: superalignment (FASTA format), partition file with gene boundaries and respective models, and the group file (NEXUS format) listing the species included in the defined groups (see Supplementary Table 13).

## Notes

### Competing Interest Statement

The authors have declared no competing interest.

